# Cross-modal interactions and movement-related tactile gating: the role of vision

**DOI:** 10.1101/2023.10.30.564759

**Authors:** Maria Casado-Palacios, Alessia Tonelli, Claudio Campus, Monica Gori

## Abstract

When engaging with the environment, multisensory cues interact and are integrated to create a coherent representation of the world around us, process that has been suggested to be affected by the lack of visual feedback in blind individuals. In addition, the presence of voluntary movement seems to be responsible for suppressing somatosensory information that is processed by the cortex, which can lead to a worse encoding of tactile information. In this work, we aim to explore how multisensory processing can be affected by active movements and the role of vision in this process. To this end, we measure the precision of 18 sighted controls and 18 blind individuals in a velocity discrimination task. The participants were instructed to detect the stimulus that contained the faster speed, between a sequence of two, in both passive and active touch conditions. The sensory stimulation could be either just tactile or audio-tactile, where a non-informative sound occurred simultaneously with the tactile stimulation. The measure of precision was obtained by computing the Just Noticeable Difference (JND) of each participant. The results show worse precision in the bimodal audio-tactile sensory stimulation in the active condition for the sighted group but not for the blind one. In the sighted group, the noise of the tactile feedback might make them more vulnerable to the noisy interference of the audio modality; however, this is not the case for the blind one, which seems to be only affected by the movement itself. Our work should be considered when developing next-generation haptic devices; moreover, it supports the need for action in the blind population.

## Introduction

When we interact with our environment, we are flooded with information from multiple sensory channels, such as when we drum our fingers on the table and produce a sound or when we move our feet in the sea and we see and hear them splash. In these moments, our brain integrates the information from the multiple senses into one unified percept to provide us with a coherent interpretation of the world. Nevertheless, when we receive multisensory information, one sense can dominate our perception due to its stronger reliability (Ernst et al., 2000; Parise & Ernst, 2017). In contrast, when we face sensory ambiguity, multisensory cues can be integrated to infer the most likely interpretation of the sensory input (Parise & Ernst, 2017). Many studies report that there is a perceptual benefit when multisensory integration is triggered. For instance, reduced reaction times have been found in the presence of redundant multisensory feedback (Pomper et al., 2014). Nonetheless, multisensory interaction can also be detrimental when conflicting information from one sensory modality affects the perception of a second modality (Keil, 2020) or acts as a distractor (Mozolic et al., 2008).

Interestingly, it has been suggested that multisensory processing is a late development achievement, as children do not show an adult-like integration until 8-10 years of age. Until this time, one sensory modality always dominates one’s perception – regardless of its reliability (Gori et al., 2008; Nardini et al., 2008). The cross-modal calibration theory states that before multisensory processing is possible, a continuous recalibration is needed in the different perceptual systems while this processing develops. During this development, cross-sensory comparison plays a crucial role; typically, people use the sense that provides the most relevant information to calibrate the others (Gori et al., 2008). This process leads to optimal integration by allowing a proper weighting of the sensory signals available (Nardini et al., 2013; Petrini et al., 2014). In the perception of space, the visual modality conveys the most reliable information (Alais & Burr, 2004). What happens when vision is missing? The study of multisensory perception in blindness allows us to understand the role of vision in this process. Previous research shows that the absence of visual input can indeed impact multisensory processing, as there is a reduced interaction between the tactile and auditory feedback in the perception of space (e.g., Hötting & Röder, 2004; Occelli et al., 2012). This likely occurs due to the lack of visual calibration over other sensory modalities. In particular, Hötting and Röder (2004) reported that congenitally blind participants were less affected by task-irrelevant stimuli than their sighted counterparts when performing a tactile-auditory illusion task. Similarly, Occelli and colleagues (2012) obtained the same results using a ventriloquist paradigm. This task consists of the biased localization of one stimulus perceived from one sensory modality by another simultaneous and spatially discrepant stimulus conveyed by another modality. These researchers discussed the reduced ventriloquist effect found in the congenitally blind group as evidence of the diminished audio-tactile interaction caused by a lack of visual input early in life (Occelli et al., 2012).

However, in multisensory research, the interaction between audio and touch is an underexplored topic, even in the sighted population. This is especially true when somatosensory feedback is generated during active touch. Active touch, which is defined as movement elicited voluntarily by the participant, is argued to be the most natural form of touch. It differs from passive tactile perception not just in terms of the mechanoreceptors activated and the processes involved (Chapman, 1994; Chapman et al., 1996) but also because the movement generated during active touch adds an obstacle to the processing of tactile inputs in the somatosensory cortex. A self-elicited movement can gate, or diminish, the transmission of tactile signals to the parietal centers involved in their decoding (Kurz et al., 2018; Williams et al., 1998). This phenomenon is known as movement-related tactile gating (e.g., Chapman, 1994; Chapman et al., 1996; Kurz et al., 2018). Surprisingly, although the importance of movement during the tactile experience has been highlighted (Cybulska-Klosowicz et al., 2011; Gibson, 1962), the impact of this phenomenon on multisensory processing, and the role of vision on it, have not yet been traced.

In this study, we explored whether multisensory processing can be affected by self-elicited movement and how vision can modulate this effect. We used the same paradigm used in previous studies (Casado-Palacios et al., 2023): using dynamic stimuli to ensure more accurate perception (Chapman et al., 1996), we developed a two-alternative-force-choice (2AFC) task to evaluate the velocity discrimination precision. Here, we present the data from a unimodal tactile that was obtained and discussed in a previous experiment (Casado-Palacios et al., 2023) and a bimodal audio-tactile sensory stimulation using passive and active touch. In the later, we included non-informative auditory feedback to accompany the tactile one. By comparing the precision in the bimodal audio-tactile sensory stimulation to the unimodal tactile in passive and active touch conditions, we aim to define how multisensory processing can be affected by active movement. In addition, including both sighted and blind participants in this study enables us to determine whether visual experience can alter this process.

We hypothesized (1) that there would be a non-significant difference between the unimodal and the bimodal stimulations for both groups in the passive condition as well as (2) in the active condition in blind participants. In contrast, we expected (3) significantly different performances between unimodal tactile and bimodal audio-tactile stimulations during active touch in the sighted participants. The results obtained supported our hypothesis.

## Methods

### Participants

This paper considered the data collected in our previous experiment (Casado-Palacios et al., 2023). The tactile conditions (passive and active) we implemented were the same data from the previous study (Casado-Palacios et al., 2022). They were used as a baseline to define the participants’ ability to perceive the speed tactile for this paper. Indeed, in the present study, we recalled previous participants to complete the audio-tactile conditions tested in the current experiment.

18 blind participants (10 women, mean age +-SD: 41.67+-11.9 years) and 18 age-matched sighted individuals (12 women; mean age +-SD: 35.11 +-11.72) took part in our study (age difference not significant: t(34)=1.67, p =0.105). Table 1 summarizes the clinical information of the blind group. All subjects stated they had no history of sensory-motor (besides blindness) or cognitive deficits. The local health service’s ethics committee (Comitato Etico, ASL3 Genovese, Italy) approved the research protocol (Comitato Etico Regione Liguria, Genoa, Italy; Prot. IIT_UVIP_COMP_2019 N. 02/2020, 4 July 2020) and the experiment was performed according to the Declaration of Helsinki. Informed consent was obtained from all participants.

**Table 1:**
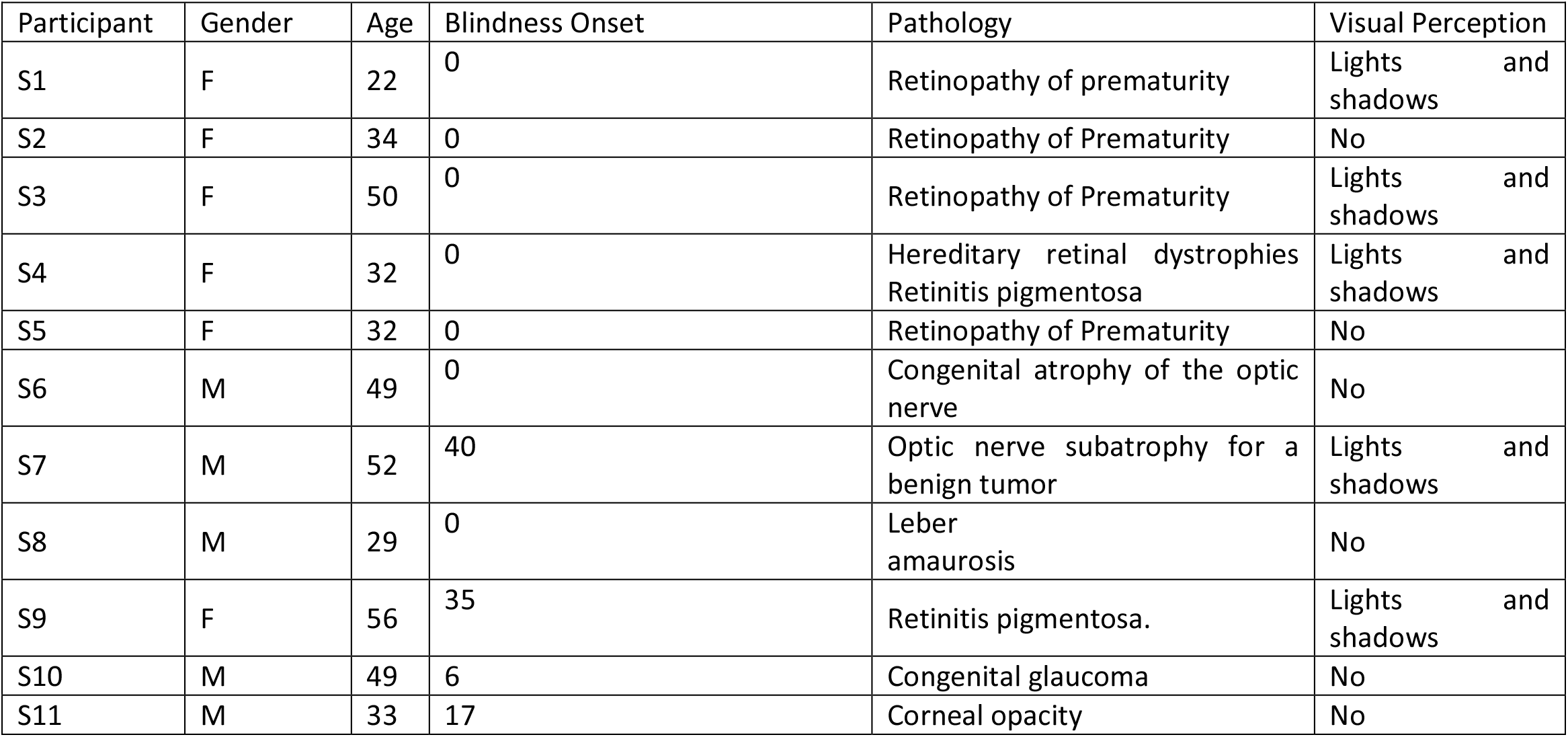
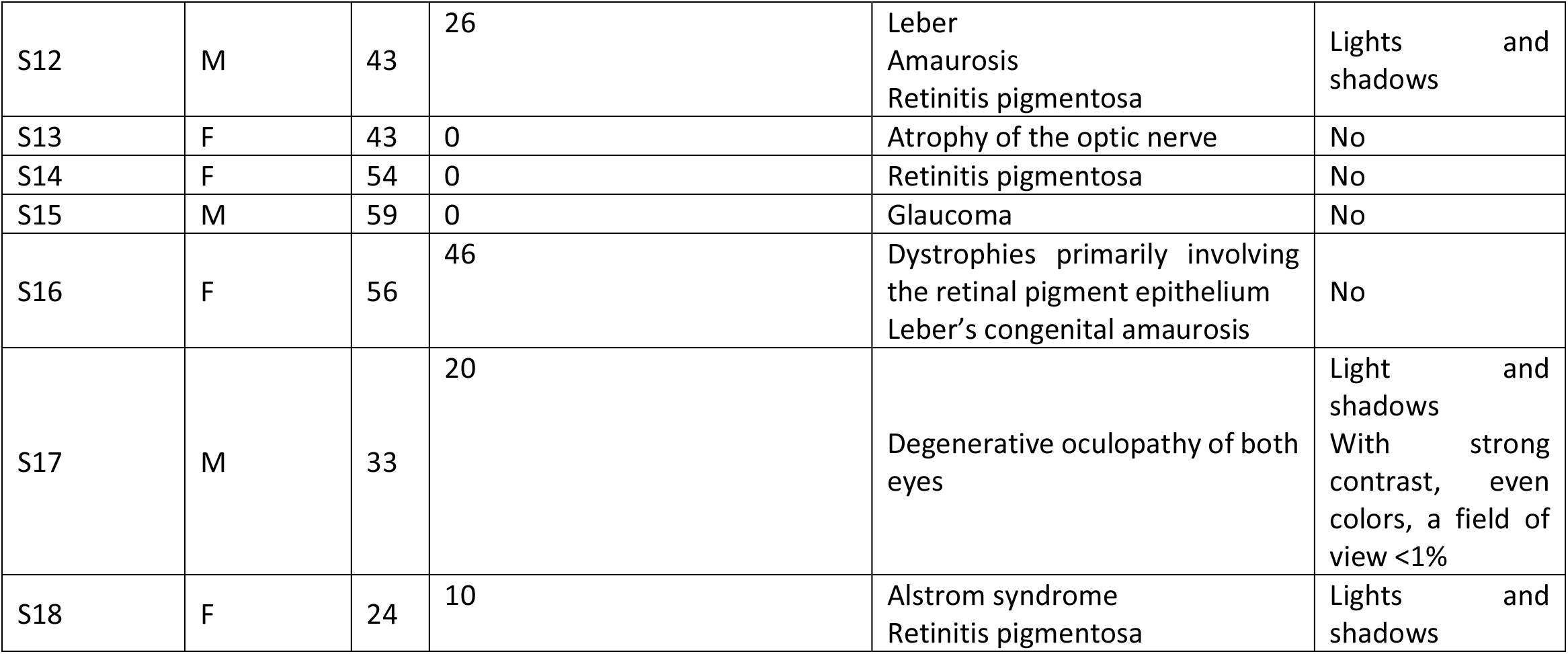
Blind participants’ clinical details, including gender, age, timing of blindness onset, pathology, and remaining visual perception.

### Design and Procedure

To induce the tactile stimulation, we used a custom-made device (Gori et al., 2011; Tomassini et al., 2012) (figure 1A). Our experiment used a wheel with 10 cycles/cm of sinusoidal greeting that could move at different speeds. The auditory stimulation was a 927Hz tone provided by a module of the MSI Caterpillar (Gori et al., 2019) attached to the wheel (figure 1B). Both devices were operated using a custom Matlab code (R2020b, The MathWorks, USA).

**Figure 1:**
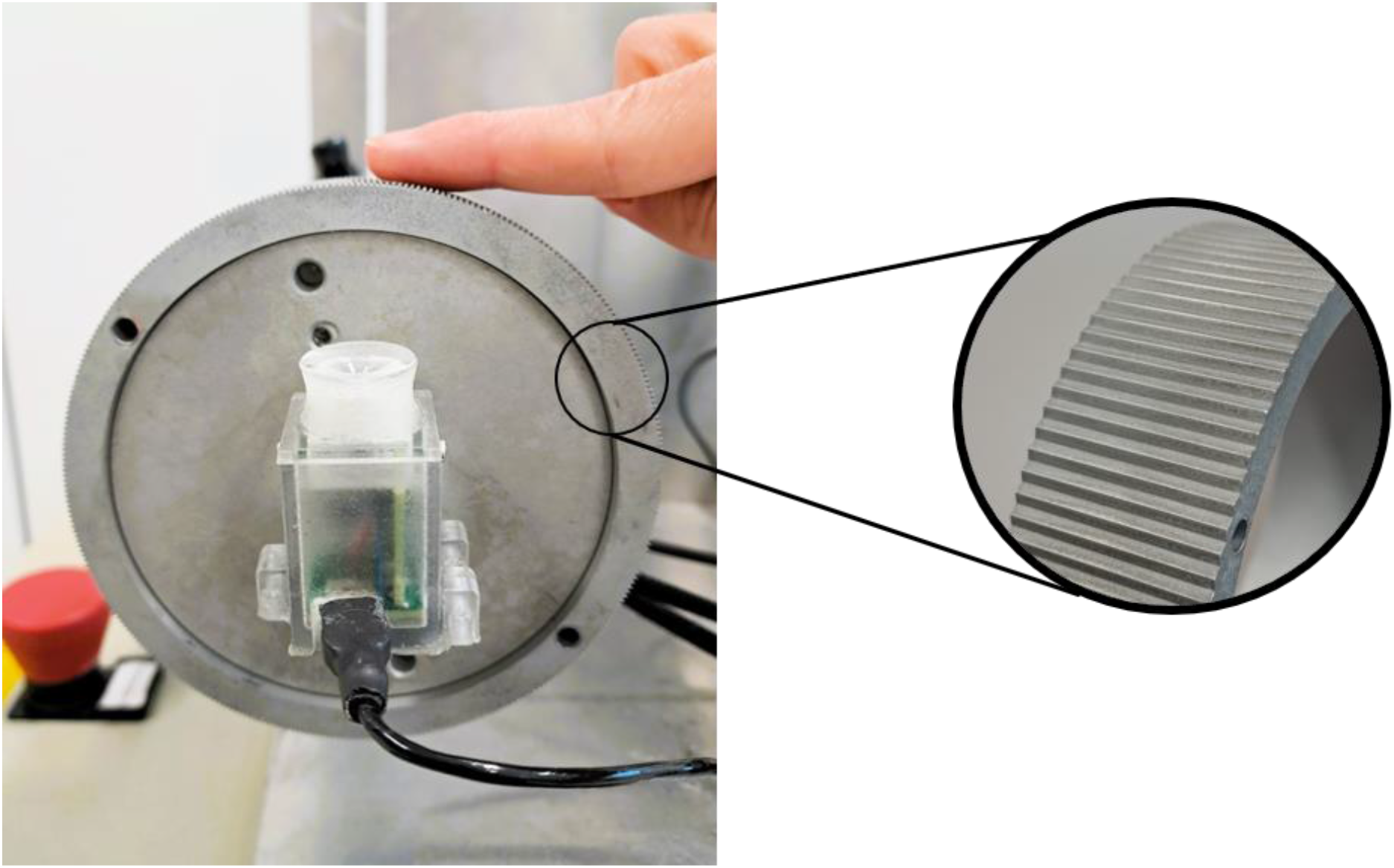
Experimental Setup consisting of a 10 cycles/cm wheel with an auditory device attached to it. The position of the finger during the experimental task is also represented.

The participants sat in a chair during the experiment with their body’s midline centered on the wheel. Figure 1 presents the wheel’s orientation and the finger’s position. Stimulation was applied to the fingertip of the index finger on the right hand. Both the blind and sighted participants were blindfolded, removing the sight variable from both groups. During the task, the participants felt two consecutive speeds lasting one second each: the first was presented at a standard velocity of 3cm/s while the second was presented at a random speed determined by the adaptive algorithm QUEST (Watson & Pelli, 1983). 500ms separated the two stimulations. The participants were required to detect the faster stimulus and report it verbally to the experimenter. We had two sensory stimulations: (1) tactile (T) and (2) audio-tactile (AT). The tactile sensory stimulus could be perceived either in (1) a passive condition, in which the finger was fixed in a specific position, or (2) an active condition, in which the finger would be actively moved in the opposite direction of the wheel’s movements by the participant. There were 30 trials per condition; thus, there were 120 in total. During the tactile-only sensory stimulus, a sound-isolating headphone was provided to eliminate the noise produced by the device to ensure the processing of only tactile information. Instructions were given to control the velocity of the finger (Togoli et al., 2020); participants were trained to move it at approximately 3cm/s.

### Data Analysis

We fitted the data using cumulative Gaussians for each condition. The best-fitting function’s mean and standard deviation yielded both the point of subject equality (PSE) and the just noticeable difference (JND) estimates. The Standard Errors for both estimates were obtained by bootstrapping (Efron & Tibshirani, 1993). In this case, we used the JND as a dependent variable.

Next, we manually inspected each psychometric curve to identify those that were inverted (i.e., negative JND) and, conservatively, we replaced the negative JND with the maximum JND value possible. In our case, it was 6 cm/s. Two blind individuals presented a negative JND. One showed an inversion in audio-tactile during active touch and the other showed inversion for the same sensory modality in passive touch. Subsequently, we evaluated the presence of outliers for all the conditions, selecting those whose JND values were lower than the first quartile minus 3 times the interquartile range as well as those greater than the third quartile and more than 3 times the interquartile range. One outlier was detected in the blind group for the audio-tactile in the passive condition. Consequently, 18 sighted and 17 blind participants were included in the analysis.

As the timing of blindness onset of our sample varied between participants, a Pearson correlation analysis was computed to determine whether to group the blind sample. The “cor.test” function from the R package “stats” (R Core Team, 2013) was applied for this analysis using the JND for each sensory stimulation (Tactile and Audio-Tactile) and condition (Passive, Active) along with the blindness duration.

The normality assumptions were tested using the Jarque Bera test (see supplementary materials): when the test resulted significant, Wilcoxon tests were adopted using “wilcox.test” function in R (package “stats”) (R Core Team, 2013) instead of t tests.

Given the results of our previous study (Casado-Palacios et al., 2023) and the research questions of the current one, we made planned comparisons between our two sensory modalities (Tactile and Audio-Tactile) separately for the passive and active touch.

To further support our results, we computed the Delta (SensoryDelta) from the bimodal (Audio-Tactile) and the unimodal (Tactile) modalities by subtracting the JND from the second to the first, and then we performed planned comparisons against zero. This final step was repeated for the Delta (ConditionDelta) from the active and passive perceptions, again by subtracting the JND from the second condition to the first one, to explore the implications of the movement itself for our two sensory conditions.

## Results

We found no correlations between the JND value and the blindness duration for the passive condition through the unimodal tactile stimulation (r= .04, t(15)= .156, p=.8785) or the bimodal audiotactile one (r= -.153, t(15)= -.598, p=.559). The same held true for the active condition for both the unimodal tactile simulation (r= -.227, t(15)= -.901, p=.382) and the audio-tactile one (r= -.108, t(15)= - .419, p=.681). As a result, we merged all the blind participants in a single group.

Worse performance emerged for audio-tactile than for the tactile sensory stimulus within the active condition for sighted individuals (Z= 0.446, p=.046) (Figure 2A). Interestingly, this difference was not present in either the passive one (Z= .046, p= .837) (Figure 2B) or the blind group (active: Z= 0.146, p= .513; passive: Z=.166, p= .459). These results suggest that only the sighted participants were negatively affected by the non-informative auditory feedback when a movement was involved.

**Figure 2:**
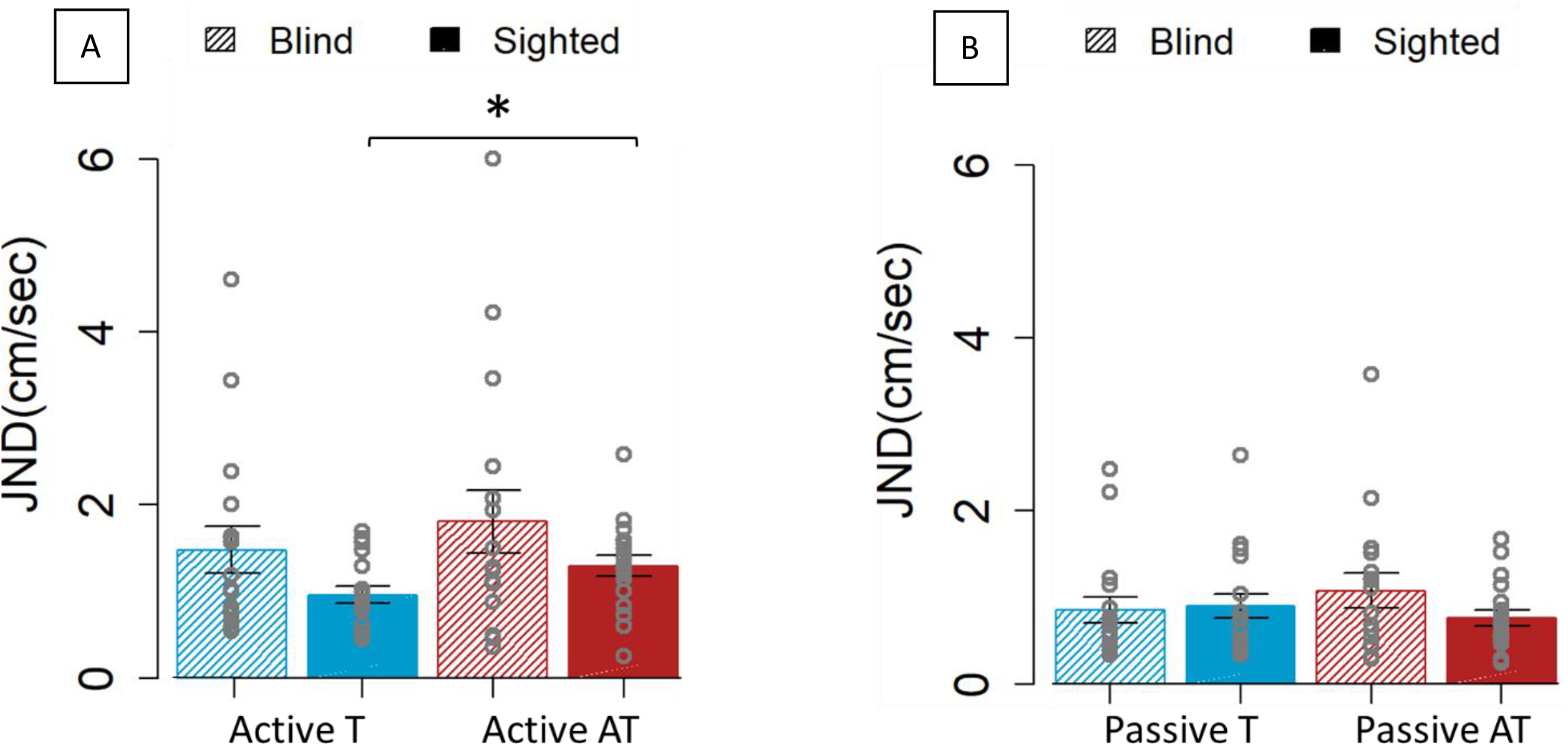
JND for the blind (45° lined boxes) and sighted individuals (full colored boxes) for the two sensory modalities. The blue color represents the tactile (T) sensory stimulus, while the red represents the audio-tactile (AT) sensory stimulus. Figure 2A shows the JND obtained in the active condition. Figure 2B illustrates the performance in the passive condition. Only the sighted participants had significantly worse precision in the presence of the non-informative sound during active movement when compared to only tactile feedback. *p<0.05.

To support our results, we computed the SensoryDelta from the audio-tactile and tactile sensory stimulations (Figure 3) for both the active and passive conditions and we compared it to zero. Here, as we expected, the only significant result was observed in the sighted group on the active condition (t(17)= 2.363, p =.03), which had a positive delta value (i.e., a higher JND in the bimodal sensory stimulation than in the unimodal one). This indicates that only the sighted participants presented a difference in the audio-tactile performance compared to the tactile one; it was significantly different from zero when a self-elicited movement was performed.

**Figure 3:**
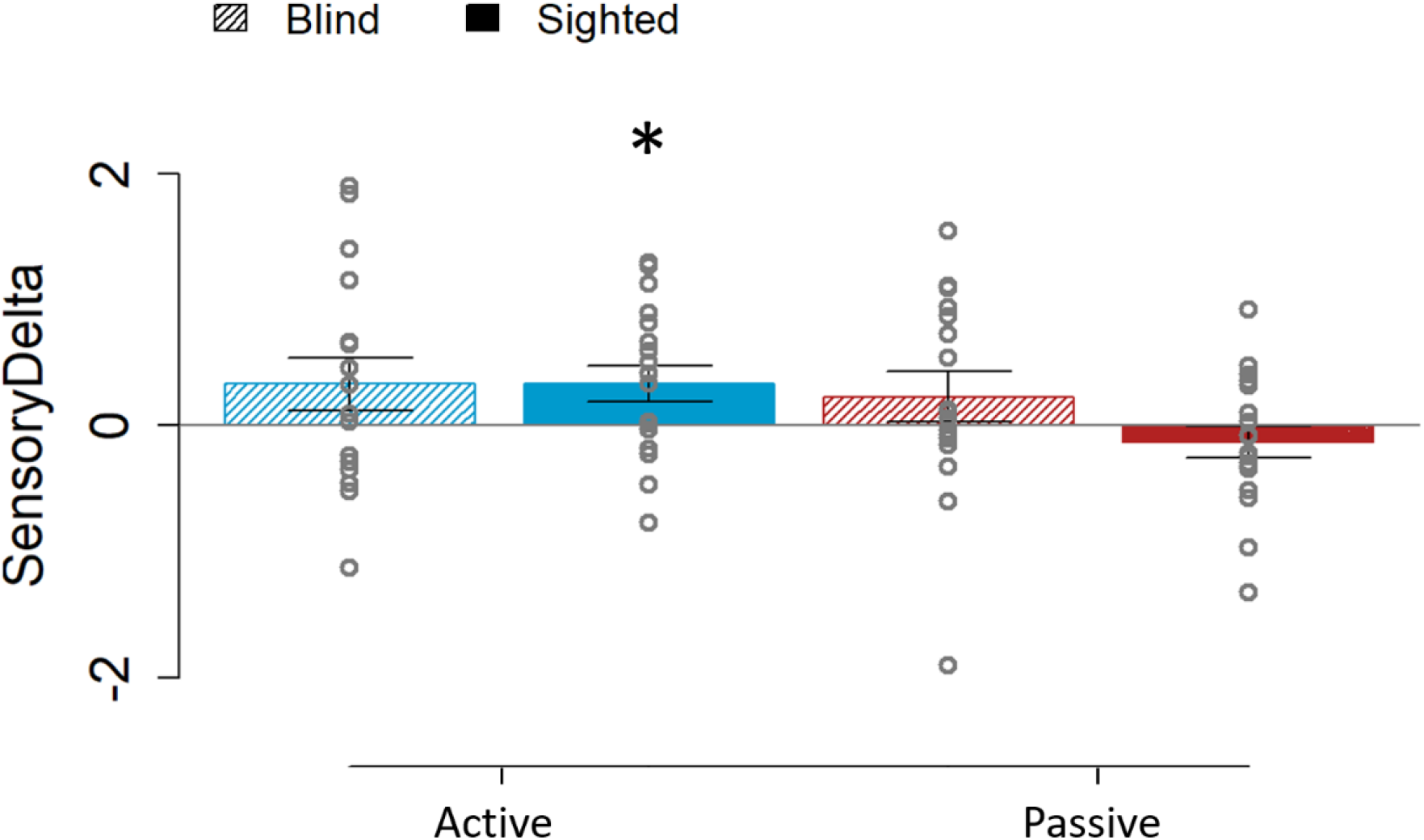
SensoryDelta values for the blind (45° lined boxes) and sighted individuals (full colored boxes) in the active (blue) and passive (red) conditions. Comparing against zero, the difference was significant for the sighted group on the active condition while it was not for the passive one or for the blind group (for any condition). *p<0.05.

Finally, to have a clearer picture of the implication of the movement in our results, we calculated the ConditionDelta from both the active and the passive conditions, for the tactile and audio-tactile sensory stimulus (Figure 4). In this case, in the blind group, there was a positive delta value (resulting from a higher JND in the active condition than the passive one) that was significantly different from zero for both the bimodal audio-tactile (t(16)= 2.804, p =.013) and unimodal tactile (t(16)= 2.599, p =.019). In contrast, in sighted individuals, only the bimodal sensory stimulus had a positive delta value significantly different from zero (audio-tactile: t(17)= 3.31, p =.004; tactile: Z= .268, p =.2309). These results implied that the difference in the precision between the active and the passive conditions is always significant in blind individuals, with higher JND in the active condition for both unimodal and bimodal sensory stimuli. However, in their sighted counterparts, the precision is only negatively affected in the active condition when the movement is coupled with non-informative feedback.

**Figure 4:**
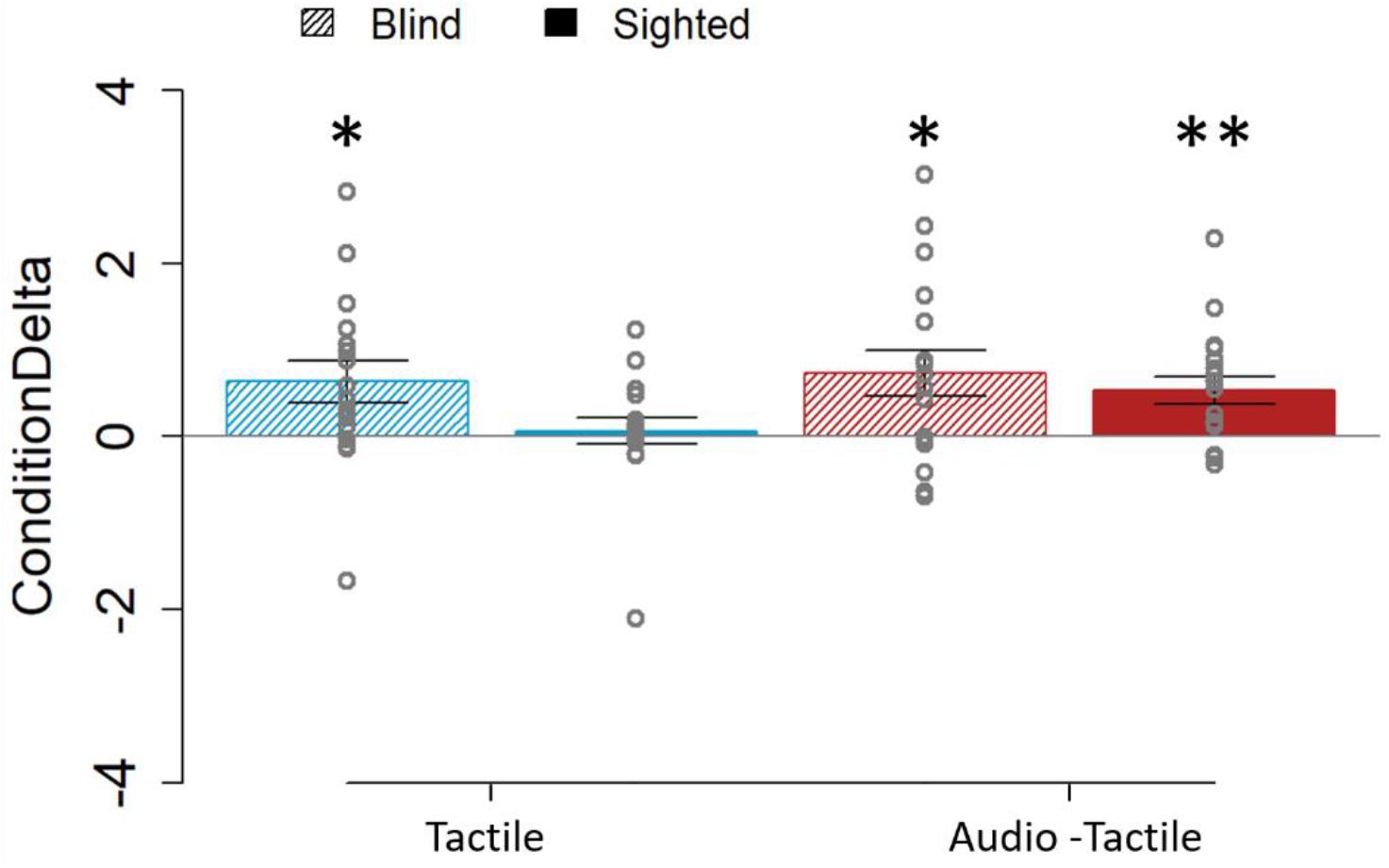
ConditionDelta values for the blind (45° lined boxes) and sighted individuals (full colored boxes) with the tactile (blue) and audio-tactile (red) stimuli. Compared to zero, the difference was significant for the blind group on the tactile and audio-tactile conditions. In the sighted group, it was only significant for the bimodal (audio-tactile) one. *p<0.05. **p<0.01.

## Discussion

When the signal from one sensory modality increases its noise, humans combine the information conveyed by different senses to create a more reliable percept of their environment (Parise & Ernst, 2017). During active touch, although we know that the movement of the body parts can affect the encoding of tactile information (Chapman et al., 1996; Fuehrer et al., 2022; Voudouris & Fiehler, 2022), researchers have neglected to study how active movement can influence multisensory processing and what the role of vision is in this process.

In this study, we found (1) a non-significant difference between the unimodal and the bimodal stimulation for both groups in the passive condition as well as (2) in the active condition for blind participants. However, there were (3) significantly different performances between unimodal tactile stimulation and the bimodal audio-tactile one during active touch in the sighted participants. In other words, the results obtained from this experiment supported our hypotheses.

We aimed to study how multisensory interaction, using tactile and auditory feedback, can be affected by movements and the modulating role of visual calibration. To do this, we provided dynamic auditory and tactile stimulation on the index fingers of the participant using a custom-made auditory device and a physical wheel. While discriminating for differences in speed, we measured the precision in sighted individuals in a pure tactile modality (the dataset was obtained from a previous experiment (Casado-Palacios et al., 2022, 2023)) and in a bimodal audio-tactile one. The tactile perception could be either passive, where participants maintained a fixed finger and felt the movement of the rotating wheel, or active, a condition in which participants moved their finger contrary to the wheel’s movement.

Our first result showed that members of the sighted group had worse precision in the presence of the bimodal audio-tactile sensory stimulus than in the tactile only one when active movements were involved but not when their perception was static. In contrast, no difference was found in the blind group. This indicates that they had similar precision for the unimodal and bimodal sensory stimulus regardless of the presence or absence of movement. To disentangle and support these results, we computed the SensoryDelta from the audio-tactile and tactile conditions (i.e. subtracting the precision obtained in the unimodal condition from the bimodal one) for passive and active touch, separately for blind and sighted participants. In line with the previous findings, there was only a significant positive SensoryDelta value for the sighted group in the active condition. This implies that for this population, the difference between the bimodal and unimodal stimulations was significantly greater than zero only in the presence of active touch. In other words, our results suggest that in the sighted group, non-informative auditory inputs only affect their precision when they perform active movements. The movement-related tactile gating might explain this effect. During active touch, the amount of information processed by the cortex is reduced (Chapman, 1994; Kurz et al., 2018); this increases the ambiguity of tactile information and makes sighted participants more vulnerable to the noise of the auditory signal.

Nevertheless, this explanation does not fully apply to the blind group. Indeed, sound’s presence did not impact the precision of this group, regardless of the type of tactile perception. This supports previous studies that extensively reported a reduced interaction between touch and audition in this population (Champoux et al., 2011; Collignon et al., 2009; Hötting & Röder, 2004; Occelli et al., 2012). A successful integration of multisensory information is required to decode between different reference frames and to know the cues that can be merged into a single perception (Nardini et al., 2008). To achieve this milestone, the cross-modal calibration theory states that the sense that offers the most robust information calibrates and tunes the rest of the sensory modalities, allowing a posterior proper weighting of the sensory signals and, thus, resulting in its integration. As vision, the most relevant sense in spatial perception, is missing in blindness, this tune between senses might fail, weakening multisensory processing and the automatic combination into a common reference frame (Collignon et al., 2009; Occelli et al., 2012; Vercillo et al., 2018) .

In addition, to dive into the implications of self-elicited movement in these results, we also calculated the ConditionDelta from the active and passive conditions for both sensory stimuli. Sighted individuals showed a significant positive difference to zero in the bimodal condition but not in the unimodal condition. This suggests that the presence of movement only caused worse precision when it was accompanied by the noise created by the non-informative sound in the control participants. In contrast, the blind group presented significant results in both sensory stimuli, always showing a worse performance in the active condition. In line with our previous work, in which we focused on the effects of self-elicited movement on tactile perception, the significance of the ConditionDelta from the active and passive touch conditions for both the tactile and audio-tactile feedback reflects that, in this population, the presence of proprioception and the predicted sensory feedback are responsible for the decreased tactile reliability (Casado-Palacios et al., 2023). This might be due to a possible inefficient integration between the cutaneous, proprioceptive, kinesthetic, and motor cues (see Casado-Palacios et al., (2023) for further details about this phenomenon). Additionally, this supports the reduced interaction between senses in this population.

## Conclusion

In our daily lives, we are exposed to multisensory cues that allow us to create a coherent representation of our environment, with which we relate through our actions. However, the use of active touch in multisensory research is limited, especially in conditions of visual deprivation. Our work shed light on this topic. Our work suggests that active touch can introduce noise to the sensory signal, modulating multisensory interaction in sighted participants, making them vulnerable to the noise interference of auditory feedback. These results highlight the need to select the proper auditory stimulation: non-informative auditory signals might interact with the tactile cues processed during movement and cause a detrimental effect on the tactile perception of this group. However, the precision of blind participants only seems affected by movement and never by non-informative auditory feedback. This finding emphasizes the need for interventions to help this population, fostering the creation of contingencies between their sensory modalities. Future work should explore whether this is also true when auditory feedback can also provide relevant information to the signal. Moreover, these results yield important implications that should be considered in developing next-generation haptic devices and in the rehabilitation field.

## Data Availability

Raw data is available on Zenodo repository (10.5281/zenodo.10008441 and 10.5281/zenodo.6581096). The code can be sent upon request. This study can be found as preprint in ……

## Acknowledgments

This work was funded by the European Union’s Horizon 2020 Research and Innovation Program under the Marie Skłodowska-Curie Grant Agreement No 860114 (MultiTOUCH).

## Author contributions

The study’s conceptualization and experimental design involved all authors. Data collection was conducted by M.C. and C.C. Data analysis was carried out by M.C. The first draft of the article was written by M.C. The final version of the article underwent review and was approved by all authors before submission.

## Additional Information

Competing interests: The authors declare no competing interests.

## Ethics declarations

The research protocol was approved by the local health service’s ethics committee (Comitato Etico, ASL3 Genovese, Italy) in line with the Declaration of Helsinki. All participants signed informed consent.

