## Supplementary material for "Cross-modal interactions and movement-related tactile gating: the role of vision"

**Normality assumptions**

To examine the Just noticeable difference (JND) variable, the Jarque Bera Test showed that the normality assumption was respected in the sighted group for the active conditions (Tactile: X-squared = 1.448, df = 2, p = .485; Audio-Tactile: X-squared = .652, df = 2, p = .722) and in the passive condition when Audio-Tactile feedback was provided (X-squared = 2.766, df = 2, p = .251); however, it was not respected for the unimodal Tactile presentation (X-squared = 10.297, df = 2, p = .006). For the blind group, the Jarque Bera Test indicated a normality assumption violation in the active touch condition (Tactile: X-squared = 10.092, df = 2, p = .006; Audio-Tactile: X-squared = 9.367, df = 2, p = .009) and the passive one (Tactile: X-squared = 11.636, df = 2, p = .003; Audio-Tactile: X-squared = 14.591, df = 2, p = .001).

Regarding the SensoryDelta variable, the Jarque Bera Test showed that the normality assumption was respected in the sighted group for both the active condition (X-squared =.601, df = 2, p = .741) and the passive one (X-squared = .324, df = 2, p = .851). It was also respected in the blind group for both the active touch condition (X-squared = .635, df = 2, p-value = 0.728) and the passive touch condition (X-squared = 3.244, df = 2, p-value = .198).

Regarding the ConditionDelta variable, the Jarque Bera Test showed that the normality assumption was respected in the sighted group for the Audio-Tactile sensory stimulation (X-squared = 2.307, df = 2, p-value = .317) but not for the Tactile one (X-squared = 358.73, df = 2, p-value < .001). In the blind group, the normality assumption was always respected (Tactile: X-squared = 1.546, df = 2, p-value = .462; Audio-Tactile: X-squared = 1.187, df = 2, p-value = .552).
